# Physical Basis of Large Microtubule Aster Growth

**DOI:** 10.1101/055939

**Authors:** Keisuke Ishihara, Kirill S. Korolev, Timothy J. Mitchison

## Abstract

Microtubule asters - radial arrays of microtubules organized by centrosomes - play a fundamental role in the spatial coordination of animal cells. The standard model of aster growth assumes a fixed number of microtubules originating from the centrosomes. However, aster morphology in this model does not scale with cell size, and we recently found evidence for non-centrosomal microtubule nucleation. Here, we combine autocatalytic nucleation and polymerization dynamics to develop a biophysical model of aster growth. Our model predicts that asters expand as traveling waves and recapitulates all major aspects of aster growth. As the nucleation rate increases, the model predicts an explosive transition from stationary to growing asters with a discontinuous jump of the growth velocity to a nonzero value. Experiments in frog egg extract confirm the main theoretical predictions. Our results suggest that asters observed in large frog and amphibian eggs are a meshwork of short, unstable microtubules maintained by autocatalytic nucleation and provide a paradigm for the assembly of robust and evolvable polymer networks.

## INTRODUCTION

Animal cells use asters, radial arrays of microtubules, to spatially organize their cytoplasm (Wilson, 1896). Specifically, astral microtubules transport organelles (Grigoriev et al., 2008; Wang et al., 2013; Waterman-Storer and Salmon, 1998), support cell motility by mediating mechanical and biochemical signals (Etienne-Manneville, 2013), and are required for proper positioning of the nucleus, the mitotic spindle, and the cleavage furrow (Field et al., 2015; Grill and Hyman, 2005; Neumüller and Knoblich, 2009; Tanimoto et al., 2016; Wilson, 1896). Within asters, individual microtubules undergo dynamic instability (Mitchison and Kirschner, 1984): They either grow (polymerize) or shrink (depolymerize) at their plus ends and stochastically transition between these two states. Collective behavior of microtubules is less well understood, and it is not clear how dynamic instability of individual microtubules controls aster growth and function.

The standard model of aster growth posits that centrosomes nucleate and anchor all microtubules at their minus ends while the plus ends polymerize outward via dynamic instability (Brinkley, 1985). As a result, aster growth is completely determined by the dynamics of individual microtubules averaged over the growing and shrinking phases. In particular, the aster either expands at a velocity given by the net growth rate of microtubules or remains stationary if microtubules are unstable and tend to depolymerize (Belmont et al., 1990; Dogterom and Leibler, 1993; Verde et al., 1992).

The standard model of aster growth is being increasingly challenged by reports of microtubules with their minus ends located far away from centrosomes (Akhmanova and Steinmetz, 2015; Keating and Borisy, 1999). Some of these microtubules may arise simply by detachment from centrosomes (Keating et al., 1997; Waterman-Storer et al., 2000) or severing of pre-existing microtubules (Roll-Mecak and McNally, 2010). However, new microtubules could also arise due to a nucleation processes independent of centrosomes (Clausen and Ribbeck, 2007; Efimov et al., 2007; Petry et al., 2013) and contribute to both aster growth and its mechanical properties. Indeed, we recently observed that centrosomal nucleation is insufficient to explain the large number of growing plus ends found in asters (Ishihara et al., 2014a). Moreover, the standard model demands a decrease in microtubule density at aster periphery, which is inconsistent with aster morphology in frog and fish embryos (Wühr et al., 2008, 2010). To resolve these inconsistencies, we proposed an autocatalytic nucleation model, where microtubules or microtubule plus ends stimulate the nucleation of new microtubules at the aster periphery (Ishihara et al., 2014a,b; Wühr et al., 2009). This mechanism generates new microtubules necessary to maintain a constant density as the aster expands. We also hypothesized that autocatalytic nucleation could effectively overcome extinction of individual microtubules, and allow rapid growth of large asters made of short, unstable microtubules. However, we did not provide a quantitative model that can be compared to the experiments or even show that the proposed mechanism is feasible.

Here, we develop a quantitative biophysical model of aster growth with autocatalytic nucleation. It predicts that asters can indeed expand even when individual microtubules turn over and disappear by depolymerization. In this regime, aster expansion is driven by the increase in the total number of microtubules, and the resulting aster is a network of short interconnected microtubules. The transition from stationary to growing asters depends on the balance between polymerization dynamics and nucleation. At this transition, our theory predicts a minimum rate at which asters grow, which we define as the gap velocity. This gap velocity arises due to the dynamic instability of microtubule polymerization and excludes a wide class of alternative models. More importantly, this mode of aster growth allows the cell to assemble asters with varying polymer densities at consistently large speeds. Using a cell-free reconstitution approach (Field et al., 2014; Nguyen et al., 2014), we perform biochemical perturbations and observe the slowing down and eventual arrest of aster growth with a substantial gap velocity at the transition. By combining theory and experiments, we provide a quantitative framework for how the cell cycle may regulate the balance between polymerization dynamics and nucleation to control aster growth. We propose that the growth of large interphase asters is an emergent property of short microtubules that constantly turnover and self-amplify.

## RESULTS

### Conceptual Model for Aster Growth based on Polymerization Dynamics and Autocatalytic Nucleation

Asters are large structures comprised of thousands of microtubules. How do the microscopic dynamics of individual microtubules determine the collective properties of asters such as their morphology and growth rate? Can asters sustain growth when individual microtubules are unstable? To address these questions, we develop a theoretical framework that integrates polymerization dynamics and autocatalytic nucleation (Fig. 1A). Our main goal is to determine the distribution of microtubules within asters and the velocity at which asters grow:

**Figure 1.**
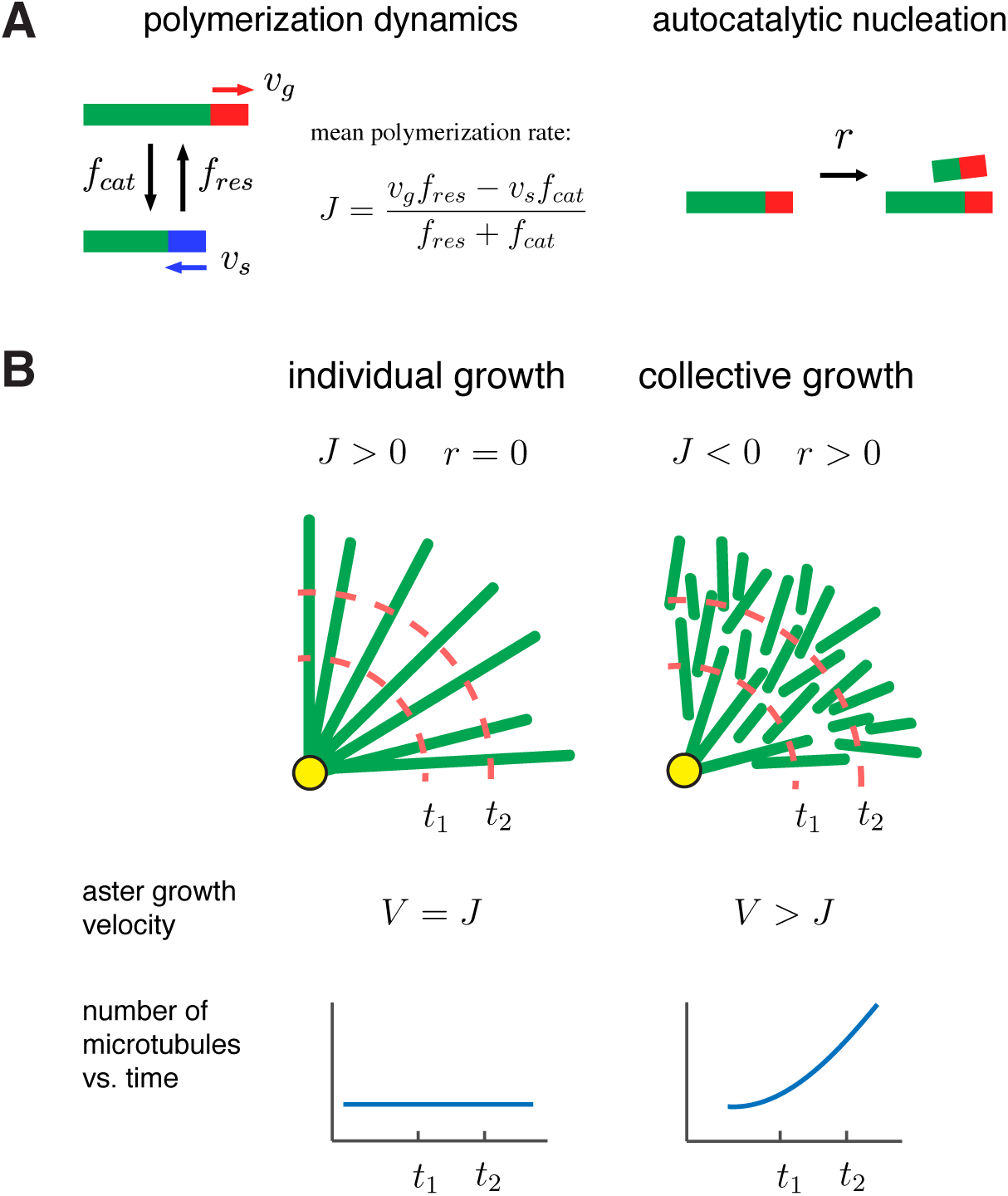
A biophysical model for the collective growth of microtubule asters. (A) We propose that asters grow via two microscopic processes: polymerization and nucleation. Individual microtubules follow the standard dynamic instability with a growing sate with polymerization rate *v*_*g*_ and a shrinking state with depolymerization rate *v*_*s*_. Transitions between the states occur at rates *f*_*cat*_ and *f*_*res*_, which model catastrophe and rescue events, respectively. New microtubules are added at a rate *r* via a nucleation at pre-existing plus ends in the growing state. (B) Individual vs. collective growth of asters. In the standard model of “individual growth”, asters increase their radius at rate 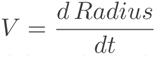 only via a net polymerization from the centrosome (yellow). Thus, this model predicts that the rate of aster growth equals the mean polymerization rate *V* = *J*, the number of microtubules is constant, and their density decreases away from the centrosomes. In the collective growth model, the microtubule density is constant and the number of microtubules increases. Autocatalytic nucleation makes asters grow faster than the net polymerization rate J and can sustain growth even when individual microtubules are unstable *J* < 0.

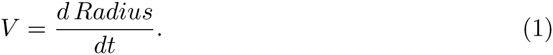

Beyond being the main experimental readout, the aster growth velocity is crucial for cell physiology because it allows large egg cells to divide its cytoplasm rapidly.

Polymerization dynamics of plus ends is an individual property of microtubules. To describe plus end dynamics, we adopt the two-state model of microtubule dynamic instability (Fig. 1A, left). In this model, a single microtubule is in one of the two states: (i) the growing state, where plus ends polymerize at rate *v*_*g*_ and (ii) the shrinking state, where plus ends depolymerize at rate *v*_*s*_. A growing microtubule may transition to a shrinking state (catastrophe event) with rate *f*_*cat*_. Similarly, the shrinking to growing transition (rescue event) occurs at rate *f*_*res*_. For large asters growing in *Xenopus* egg cytoplasm, we provide estimates of these parameters in Table 1.

**Table 1.** Model parameters used to describe large aster growth reconstituted in interphase *Xenopus* egg extract.

| Quantity | Symbol | Value | Comment |
| --- | --- | --- | --- |
| Polymerization rate | $v_g$ | 30 $\mu\text{m}/\text{min}$ | Measured from growing plus ends and EB1 comets |
| Depolymerization rate | $v_s$ | 42 $\mu\text{m}/\text{min}$ | Measured from shrinking plus ends (Ishihara et al., 2014a) |
| Catastrophe rate | $f_{cat}$ | 3.3 $\text{min}^{-1}$ | Measured from EB1 comet lifetimes (see Methods) |
| Rescue rate | $f_{res}$ | $2.0 \pm 0.3 \text{ min}^{-1}$ | Estimated from Eq. (4) and (6) |
| Autocatalytic nucleation rate | $r$ | $2.1 \pm 0.2 \text{ min}^{-1}$ | Estimated from Eq. (4) and (6) |
| Carrying capacity of growing ends | $K$ | 0.4 $\mu\text{m}^{-2}$ | Estimated from comparing $C_g^{bulk}$ to predicted (see SI) |
| Mean microtubule length | $\langle l \rangle$ | $16 \pm 2 \mu\text{m}$ | Estimated from from dynamics parameters (see SI) |
| Aster growth velocity | $V$ | $22.3 \pm 2.6 \mu\text{m}/\text{min}$ | Measured from rate of aster radius increase |
| Gap velocity | $V_{gap}$ | $12.8 \pm 1.7 \mu\text{m}/\text{min}$ | Measured from aster growth at 320 nM MCAK-Q710 |
| Bulk growing plus end density | $C_g^{bulk}$ | $0.053 \pm 0.030 \mu\text{m}^{-2}$ | Measured from EB1 comet density (Ishihara et al., 2014a) |

Plus end dynamics can be conveniently summarized by the time-weighted average of the polymerization and depolymerization rates (Dogterom and Leibler, 1993; Verde et al., 1992):

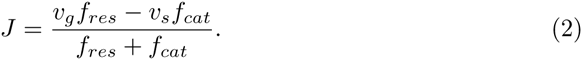

This parameter describes the tendency of microtubules to grow or shrink. When *J* < 0, microtubules are said to be in the bounded regime because their length inevitably shrinks to zero, i.e. microtubule disappears. When *J* > 0, microtubules are said to be in the unbounded regime, because they have a nonzero probability to become infinitely long. Parameter *J* also determines the mean elongation rate of a very long microtubule that persists over many cycles of catastrophe and rescue. The dynamics of short microtubules, however, depends on their length and initial state (growing vs. shrinking) and should be analyzed carefully.

The standard model posits that asters are produced by the expansion of individual microtubules, so the transition from small mitotic asters to large interphase asters is driven by a change in the sign of *J* (Dogterom and Leibler, 1993; Verde et al., 1992) (Fig. 1B left, “individual growth”). With bounded dynamics *J* < 0, the standard model predicts that every microtubule shrinks to zero length and disappears. This microtubule loss is balanced by nucleation of new microtubules at the centrosomes, the only place where nucleation is allowed in the standard model. As a result, asters remain in the stationary state and are composed of a few short microtubules, and the aster growth velocity is thus *V* = 0. With unbounded dynamics *J* > 0, the standard model predicts an aster that has a constant number of microtubules and increases its radius at a rate equal to the elongation rate of microtubules (i.e. *V* = *J*).

Below, we add autocatalytic microtubule nucleation (Fig. 1A, right) to the standard model and propose the regime of “collective growth” (Fig. 1B, right). In this regime, asters grow (*V* > 0) although individual microtubules are bounded (*J* < 0) and are, therefore, destined to shrink and disappear. The growth occurs because more microtubules are nucleated than lost, and new microtubules are typically nucleated further along the expansion direction. Indeed, when a new microtubule is nucleated, it is in a growing state and starts expanding outward before its inevitable collapse. During its lifetime, this microtubule can nucleate a few more microtubules all of which are located further along the expansion direction. As we show below, this self-amplifying propagation of microtubules is possible only for sufficiently high nucleation rates necessary to overcome microtubule loss and sustain collective growth.

Specifically, we assume that new microtubules nucleate at locations away from centrosomes at rate *Q*. This rate could depend on the local density of growing plus ends if they serve as nucleation sites or the local polymer density if nucleation can occur throughout a microtubule. This rate could depend on the local density of growing plus ends, if they serve as nucleation sites or the local polymer density if nucleation occurs along the side of pre-existing microtubules. The new microtubules have zero length and tend to grow radially due to mechanical interactions with the existing microtubule network. These non-centrosomal microtubules disappear when they shrink back to their minus ends. Our assumptions are broadly consistent with known microtubule physiology (Clausen and Ribbeck, 2007; Petry et al., 2013), and we found strong evidence for nucleation away from centrosomes in egg extract by microtubule counting in growing asters (Ishihara et al., 2014a). Below, we consider plus-end-stimulated nucleation and the analysis for the polymer-stimulated nucleation is summarized in the SI.

Without negative feedback, autocatalytic processes lead to exponential growth, but there are several lines of evidence for an apparent “carrying capacity” of microtubules in a given cytoplasmic volume (Clausen and Ribbeck, 2007; Ishihara et al., 2014a; Petry et al., 2013). Saturation is inevitable since the building blocks of microtubules are present at a fixed concentration. In our model, we impose a carrying capacity by expressing autocatalytic nucleation as a logistic function of the local density of growing plus ends, which is qualitatively consistent with local depletion of nucleation factors such as the gamma-tubulin ring complex. Other forms of negative feedback (e.g. at the level of polymerization dynamics) are possible as well. In SI, we show that the type of negative feedback does not affect the rate of aster growth, which is determined entirely by the dynamics at the leading edge of a growing aster where the microtubule density is small and negative feedback can be neglected.

### Mathematical Model of Autocatalytic Growth of Asters

Assuming large number of microtubules, we focus on the mean-field or deterministic dynamics (SI) and formalize our model as a set of partial differential equations. Specifically, we let *ρ*_*g*_(*t*; *x*, *l*) and *ρ*_*s*_(*t*; *x*, *l*) denote respectively the number of growing and shrinking microtubules of length *l* with their minus ends at distance *x* > 0 from the centrosome. The number of newly nucleated microtubules is given by *Q*(*x*) = *rC*_*g*_(*t*; *x*)(1 − *C*_*g*_(*t*, *x*)/*K*), where *r* is the nucleation rate, *K* is the carrying capacity controlling the maximal plus end density, and *C*_*g*_(*t*, *x*) is the local density of the growing plus ends at point *x*. The polymerization dynamics and nucleation are then described by,

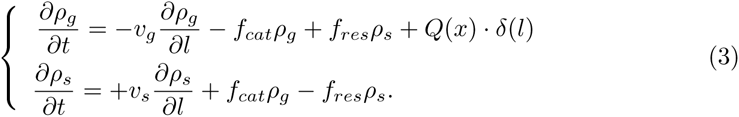

Note that polymerization and depolymerization changes the microtubule length *l*, but not the minus end position *x*. Equations at different *x* are nevertheless coupled due to the nucleation term, which depends on *x* through *C*_*g*_.

To understand this system of equations, consider the limit of no nucleation (*r* = 0). Then, the equations at different *x* become independent and we recover the standard model that reduces aster growth to the growth of individual microtubules (Dogterom and Leibler, 1993; Verde et al., 1992). With nucleation, aster growth is a collective phenomenon because microtubules of varying length and minus end positions contribute to *C*_*g*_(*t*, *x*), which can be expressed as a convolution of *ρ*_*g*_ (see SI). The delta-function *δ*(*l*) ensures that newly nucleated microtubules have zero length.

Finally, we need to specify what happens when microtubules shrink to zero length. In our model, microtubules originating from centrosomes rapidly switch from shrinking to growth (i.e. re-nucleate), while non-centrosomal microtubules disappears completely (i.e. no re-nucleation occurs). We further assume that mother and daughter microtubules disappear without affecting each other. Indeed, if the collapse of the mother microtubule triggered the collapse of the daughter microtubule (or vice versa), then no net increase in the number of microtubules would be possible in the bounded regime. One consequence of this assumption is that the minus end of a daughter microtubule becomes detached from any other microtubules in the aster following the collapse of the mother microtubule. As a result, minus ends need to be stabilized after nucleation possibly by some additional factors (Akhmanova and Hoogenraad, 2015) and mechanical integrity of the aster should rely on microtubule bundling (Ishihara et al., 2014a).

### Asters Can Grow as Spatially Propagating Waves with Constant Bulk Density

To check if our model can describe aster growth, we solved Eq. (3) numerically using finite difference methods in an 1D planar geometry. With relatively low nucleation rates and *J* < 0, microtubule populations reached a steady-state profile confined near the centrosome reminiscent of an aster in the standard model with bounded microtubule dynamics (Fig. 2A left). When the nucleation rate was increased, the microtubule populations expanded as a travelling wave with an approximately invariant shape and constant microtubule density at the periphery (Fig. 2A right) consistent with the growth of interphase asters in our reconstitution experiments (Fig. 2B and (Ishihara et al., 2014a)). Thus, our model predicted two qualitatively different states: stationary and growing asters.

**Figure 2.**
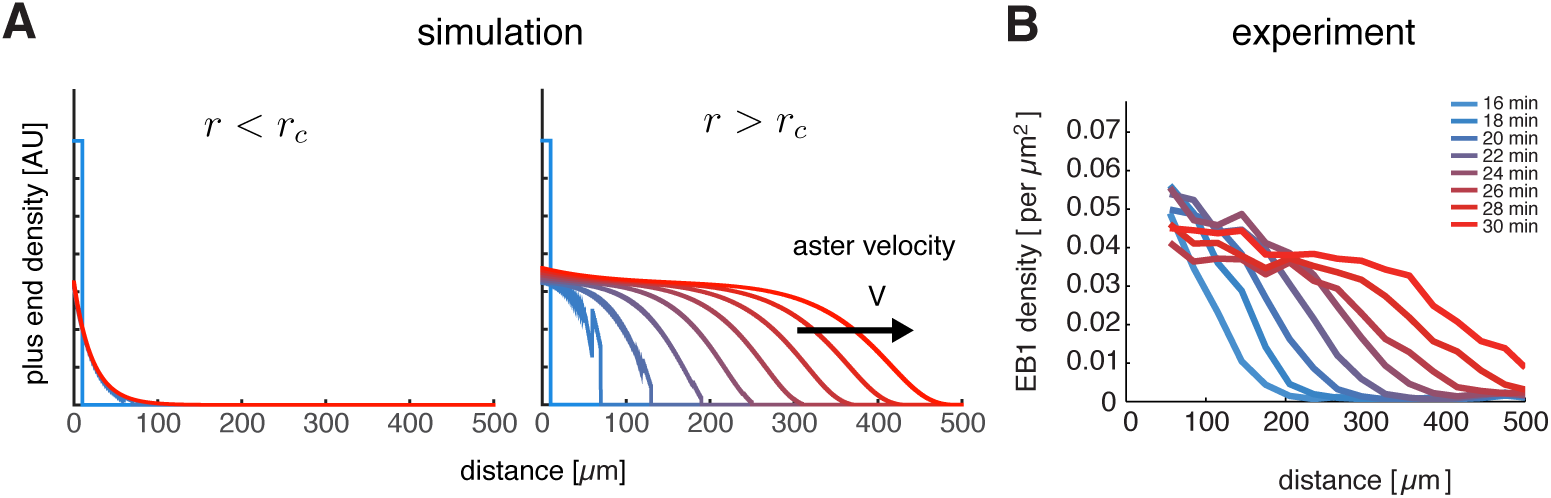
Our model captures key features of large aster growth. (A) Time evolution of growing plus end density predicted by our model, which we solved via numerical simulations in 1D geometry. In the stationary regime, the microtubule population remained near the centrosome *v*_*g*_ = 30, *v*_*s*_ = 40, *f*_*cat*_ = 3, *f*_*res*_ = 1, and *r* = 1.0 (left). In contrast, outward expansion of the microtubule population was observed when the nucleation rate was increased to *r* = 2.5, above the critical nucleation rate *r*_*c*_ (right). For both simulations, microtubules are in the bounded regime *J* < 0. (B) Experimental measurements confirm that asters expand at a constant rate with time-invariant profiles of the plus end density, as predicted by our model. The plus end densities were estimated as EB1 comet density during aster growth as previously Figdescribed.3. Reprinted with permission from (Ishihara et al., 2014a).

### Analytical Solution for Growth Velocity and Critical Nucleation

Next, we solved Eq. (3) exactly and obtained the following analytical expression for the growth velocity of an aster in terms of model parameters:

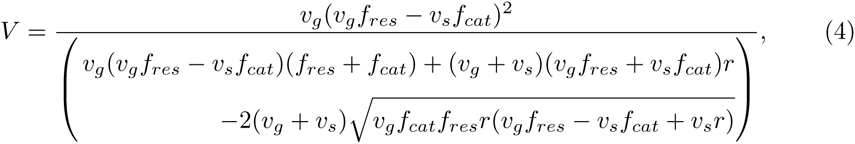

which holds for the parameter range *r*_*c*_ < *r* < *f*_*cat*_. The details of the calculation, including the definition of *r*_*c*_ are summarized in SI.

Using this expression, we summarize how aster growth velocity *V* is affected by the mean polymerization rate *J* (Fig. 3A) and nucleation rate *r* (Fig. 3B). In the absence of autocatalytic nucleation (*r* = 0), our model reduces to the standard model and predicts that asters only grow when *J* > 0 with *V* = *J* (Fig. 3A blue line). When nucleation is allowed (*r* > 0), the growth velocity increases with *r* and asters can grow even when individual microtubules are unstable *J* < 0 (Fig. 3A and 3B). During this collective growth, the aster expands radially because more microtubules are nucleated than lost at the front. In the aster bulk, nucleation is reduced from the carrying capacity, and the aster exists in the dynamic balance between microtubule gain due to nucleation and loss due to depolymerization. Since microtubules are in the bounded regime, their lifetime is short, and they disappear before reaching appreciable length. In sharp contrast to the standard model, we predict that asters are a dynamic network of short microtubules with properties independent from the distance to the centrosome. Thus, nucleation not only increases the number of microtubules, but also controls the growth rate and spatial organization of asters enabling them to span length scales far exceeding the length of an individual microtubule.

**Figure 3.**
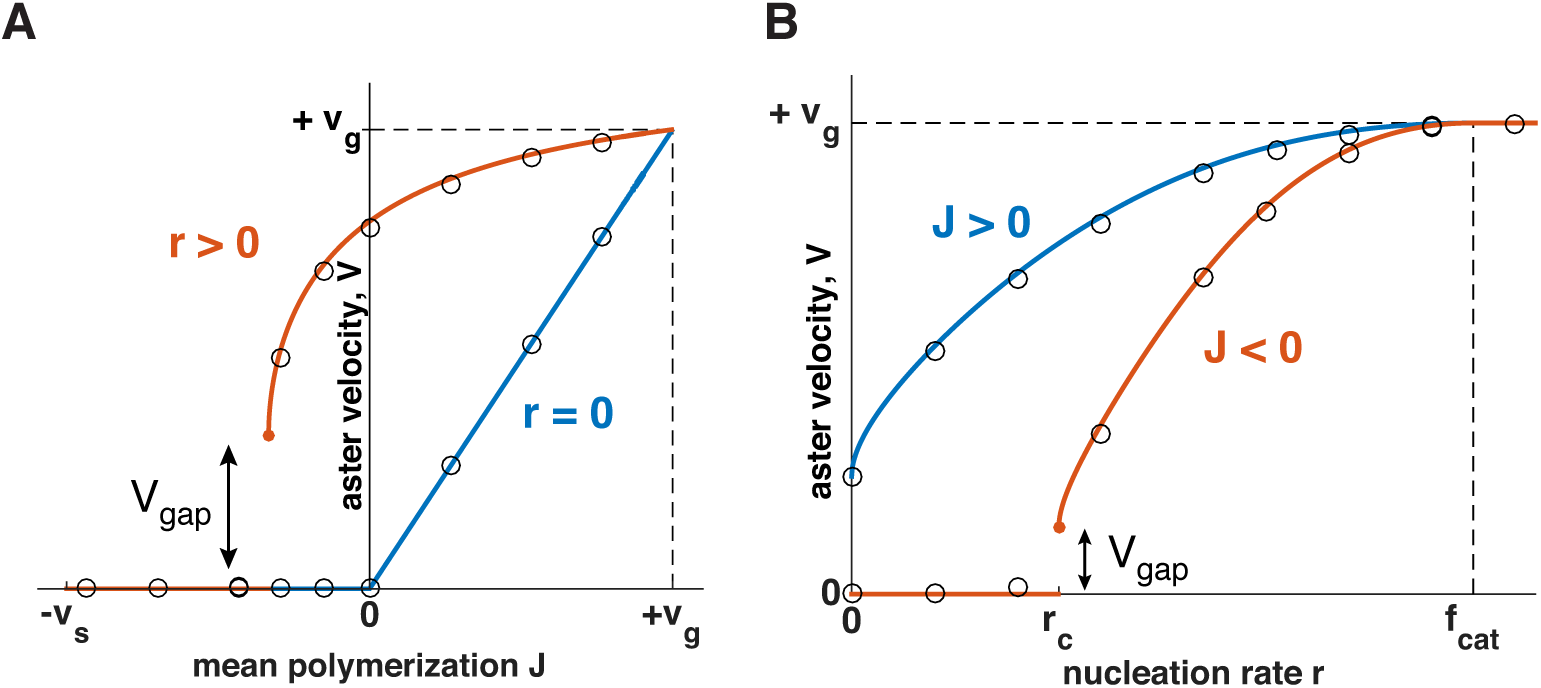
Explosive transition from stationary to growing asters and other theoretical predictions. Analytical solution (lines) and numerical simulations (dots) predict that asters either remain stationary or expand at a constant velocity, which increases with the net polymerization rate *J* (A) and nucleation rate *r* (B). The transition to a growing state is accompanied by finite jump in the expansion velocity labeled as *V*_*gap*_. (A) The behavior in the standard model (*r* = 0) is shown in blue and our model (*r* = 1.5) in red. Note that aster growth commences at *J* < 0 in the presence of nucleation and occurs at a minimal velocity *V*_*gap*_. Although spatial growth can occur for both *J* > 0 and *J* < 0 the properties of the resulting asters could be very different (see SI). Here, *v*_*g*_ = 30; *v*_*s*_ = 30; *f*_*cat*_ = 3. (B) If *J* < 0, critical nucleation *r*_*c*_ is required to commence aster growth. Blue line corresponds to *J* > 0(*v*_*g*_ = 30; *v*_*s*_ = 15; *f*_*cat*_ = 3; *f*_*res*_ = 3) and red line to *J* < 0(*v*_*g*_ = 30; *v*_*s*_ = 15; *f*_*cat*_ = 3; *f*_*res*_ = 1). See Materials and Methods and SI for the details of the analytical solution and numerical simulations.

When *J* < 0, a critical nucleation rate is required for aster growth (Fig. 3B). Indeed, microtubules constantly disappear as their length shrinks to zero, and the nucleation of new microtubules need to occur frequently enough to overcome the microtubule loss. Consistent with this argument, our analytical solution predicts no aster growth below a certain value of nucleation (SI), termed critical nucleation rate *r*_*c*_:

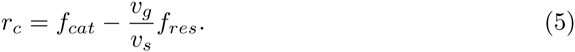

The right hand side of this equation is the inverse of the average time that a microtubule spends in the growing state before shrinking to zero-length and disappearing (SI). Thus, aster growth requires that, on average, a microtubule to nucleate at least one new microtubule during its lifetime.

The dependence of the critical nucleation rate on model parameters is very intuitive. Increasing the parameters in favor of polymerization (*v*_*g*_ and *f*_*res*_), lowers the threshold level of nucleation required for aster growth, while increasing the parameters in favor of depolymerization (*v*_*s*_ and *f*_*cat*_) has the opposite effect. We also find that *r*_*c*_ = 0 when J = 0, suggesting that there is no critical nucleation rate for *J* ≥ 0. This limit is consistent with the standard model with *J* > 0 and *r* = 0 where the aster radius increases albeit with radial dilution of microtubule density (Fig. 1B). The critical nucleation rate conveys the main implication of our theory: the balance between polymerization dynamics and autocatalytic nucleation defines the quantitative condition for continuous aster growth.

### Explosive Transition to Growth with a “Gap Velocity”

At the critical nucleation rate *r* = *r*_*c*_, the aster growth velocity *V* takes a positive, nonzero value (Fig. 3), which we refer to as the “gap velocity” (SI):

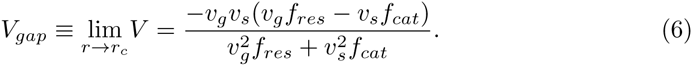

This finite jump in the aster velocity is a consequence of microtubules with finite length undergoing dynamic instability and is in sharp contrast to the behavior of reaction-diffusion systems, where travelling fronts typically become infinitesimally slow before ceasing to propagate (Chang and Ferrell, 2013; Hallatschek and Korolev, 2009; Méndez et al., 2007; van Saarloos, 2003). One can understand the origin of *V*_*gap*_ > 0 when microtubules are eliminated after a catastrophe event (*f*_*res*_ = 0; *J* = −*v*_*s*_). In this limit, plus ends always expand with the velocity *v*_*g*_ until they eventually collapse. Below *r*_*c*_, this forward expansion of plus ends fails to produce aster growth because the number of plus ends declines on average. Right above *r*_*c*_, the number of plus ends is stable and aster grows at the same velocity as individual microtubules. Indeed, Eq. (6) predicts that *V*_*gap*_ = *v*_*g*_ when *f*_*res*_ = 0. The dynamics are similar for *f*_*res*_ > 0. At the transition, nucleation stabilizes a subpopulation of microtubules expanding forward, and their average velocity sets the value of *V*_*gap*_. We also find that the magnitude of *V*_*gap*_ is inversely proportional to the mean length of microtubules in the system (SI). Thus, the shorter the microtubules, the more explosive this transition becomes.

In the SI, we also show that microtubule density inside the aster is proportional to *r* − *r*_*c*_. Thus, the density is close to zero during the transition from stationary to growing asters, but quickly increases as the nucleation rate becomes larger. As a result, cells can achieve rapid aster growth while keeping the density of the resulting microtubule network sufficiently low. The low density might be beneficial because of its mechanical properties or because it simply requires less tubulin to produce and energy to maintain. In addition, the explosive transition to growth with *V*_*gap*_ > 0 allows the cell to independently control the aster density and growth speed.

Model parameters other than the nucleation rate can also be tuned to transition asters from growth to no growth regimes. Similar to Eq. (5) and (6), one can define the critical parameter value and gap velocity to encompass all such transitions (Table S1). In all cases, we find that the onset of aster growth is accompanied by discontinuous increase in the growth velocity. The finite jump in aster growth velocity is similarly predicted in a wide range of alternative scenarios including (i) feedback regulation of plus end dynamics (SI and Figure 3 - figure supplement 1) and (ii) aster growth by microtubule polymer-stimulated nucleation (SI and Figure 3 - figure supplement 2). In summary, the gap velocity is a general prediction of the collective behavior of microtubules that are short-lived.

### Titration of MCAK Slows then Arrests Aster Growth with Evidence for a Gap Velocity

Based on our theory, we reasoned that it would be possible to transform a growing interphase aster to a small, stationary aster by tuning polymerization dynamics and/or nucleation via biochemical perturbations in *Xenopus* egg extract. To this end, we performed reconstitution experiments in undiluted interphase cytoplasm supplied with anti-Aurora kinase A antibody coated beads, which nucleate microtubules and initiate aster growth under the coverslip (Field et al., 2014; Ishihara et al., 2014a). We explored perturbation of various regulators for plus end dynamics and nucleation. We settled on perturbation of MCAK/KIF2C, classically characterized as the main catastrophe-promoting factor in the extract system (Kinoshita et al., 2001; Walczak et al., 1996), and imaged aster growth.

In control reactions, aster radius, visualized by the plus end marker EB1-mApple, increased at velocities of 20.3 ± 3.1 *μm*/*min* (n=21 asters). We saw no detectable changes to aster growth with addition of the wild type MCAK protein. In contrast, addition of MCAK-Q710 (Moore and Wordeman, 2004) decreased aster growth velocity (Fig. 4A and B). At concentrations of MCAK-Q710 above 320 nM, most asters had small radii with very few microtubules growing from the Aurora A beads. In our model, this behavior is consistent with any change of parameter(s) that reduces the aster growth velocity (Eq. (4)) and arrests growth.

**Figure 4.**
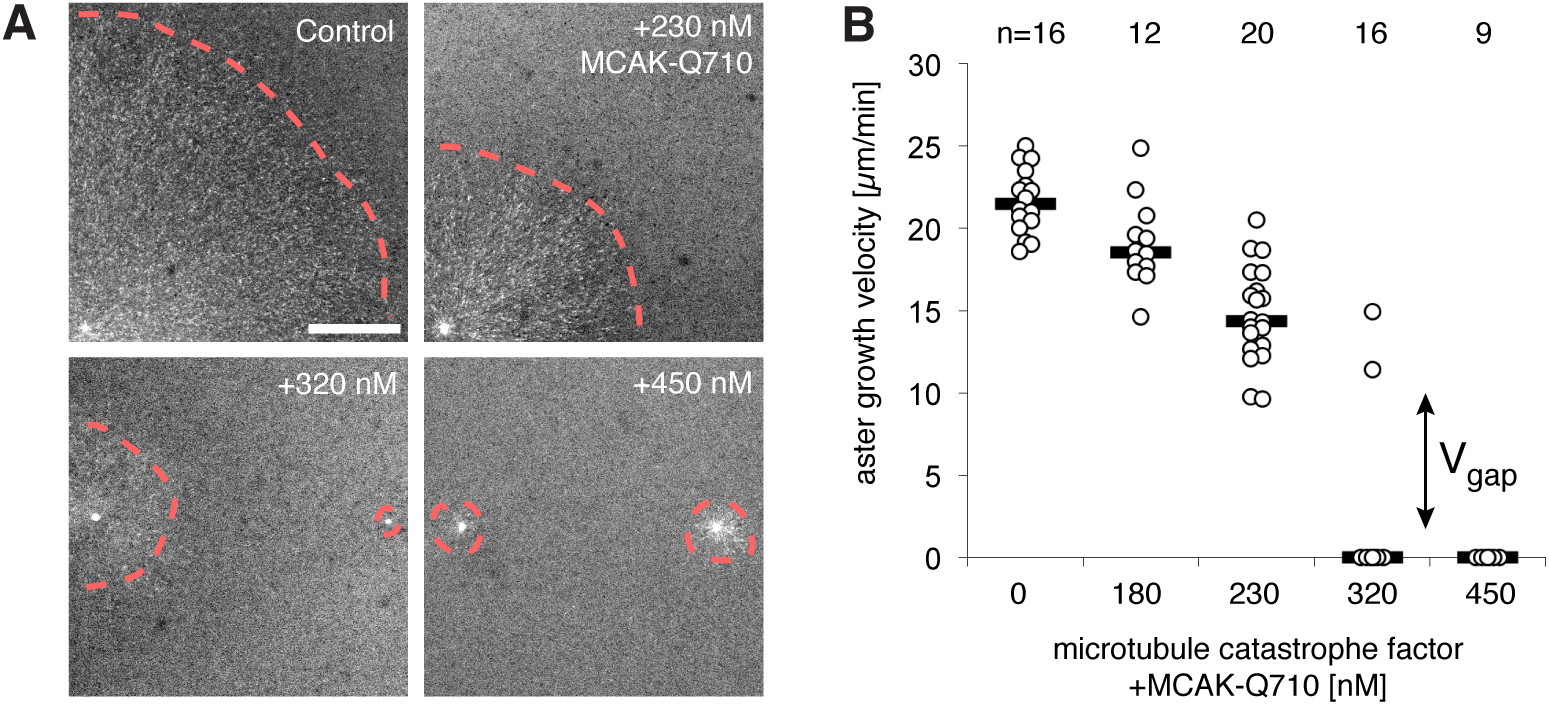
Titration of MCAK-Q710 slows then arrests aster growth through a discontinuous transition. (A) Addition of MCAK-Q710 results in smaller interphase asters assembled by Aurora A beads in *Xenopus* egg extract. Images were obtained 20 minutes post initiation with the plus end marker EB1-mApple. Dotted lines indicate the approximate outline of asters. (B) Aster growth velocity decreases with MCAK-Q710 concentration and then abruptly vanishes as predicted by the model. Note a clear gap in the values of the observed velocities and bimodality near the transition, which support the existence of *V*_*gap*_. Quantification methods are described in methods and Figure 4-figure supplement 1

At 320nM MCAK-Q710 concentration, we observed bimodal behavior. Some asters increased in radius at moderate rates, while other asters maintained a stable size before disappearing, presumably due to the decrease of centrosomal nucleation over time (Figure 4-figure supplement 1 and (Ishihara et al., 2014a)). In particular, we observed no asters growing at velocities between 0 and 9 *μm*/*min* (Fig. 4B and Figure 4-figure supplement 1). This gap in the range of possible velocities is consistent with the theoretical prediction that growing asters expand above a minimal rate *V*_*gap*_.

To confirm that the failure of aster growth at high concentrations of MCAK-Q710 is caused by the changes in aster growth rather than nucleation from the beads, we repeated the experiments with *Tetrahymena* pellicles as the initiating centers instead of Aurora A beads. Pellicles are pre-loaded with a high density of microtubule nucleating sites, and are capable of assembling large interphase asters (Ishihara et al., 2014a). We found pellicle initiated asters to exhibit a similar critical concentration of MCAK-Q710 compared to Aurora A bead asters. While the majority of Aurora A beads subjected to the highest concentration of MCAK-Q710 lost growing microtubules over time, a significant number of microtubules persisted on pellicles even after 60 min (Figure 4-figure supplement 2). The radii of these asters did not change, consistent with our prediction of stationary asters. Thus, the pellicle experiments confirmed our main experimental result of small, stationary asters and that the nature of transition is consistent with the existence of a gap velocity.

Finally, we asked which parameters in our model were altered in the MCAK-Q710 perturbation. To this end, we measured the polymerization and catastrophe rates in interphase asters assembled by Aurora A beads at various MCAK-Q710 concentrations. We imaged EB1 comets at high spatiotemporal resolution, and analyzed their trajectories by tracking-based image analysis ((Applegate et al., 2011; Matov et al., 2010), Methods). Neither the polymerization nor the catastrophe rate changed at the MCAK-Q710 concentrations corresponding to the transition between growing and stationary asters (Figure 4-figure supplement 3). MCAK-Q710 has been reported to reduce microtubule polymer levels in cells (Moore and Wordeman, 2004), but its precise effect on polymerization dynamics and/or nucleation remains unknown. Our data is consistent with the following three scenarios for how MCAK-Q710 antagonizes microtubule assembly: (i) increased depolymerization rate, (ii) decreased rescue rate, and/or (iii) decreased nucleation rate.

## DISCUSSION

### An Autocatalytic Model of Aster Growth

It has not been clear whether the standard model of aster growth can explain the morphology of asters observed in all animal cells, including those of extreme size (Mitchison et al., 2015). To resolve this question, we constructed a biophysical framework that incorporates microtubule polymerization dynamics and autocatalytic nucleation. Numerical simulations and analytical solutions (Fig. 2, 3, and Fig. 3-figure supplement 1, 2) recapitulated both stationary and continuously growing asters in a parameter-dependent manner. Interestingly, the explosive transition from “growth” to “no growth” was predicted to involve a finite growth velocity, which we confirmed in biochemical experiments (Fig. 4).

Our biophysical model offers a biologically appealing explanation to aster growth and allows us to estimate parameters that are not directly accessible: the rescue and autocatalytic nucleation rates. For example, if we assume that MCAK-Q710 decreases the nucleation rate, we may use the *V*_*gap*_ equation for *r* → *r*_*c*_ (Eq. (6)), the equation for aster growth velocity V (Eq. (4)), and our measurements of *v*_*g*_, *v*_*s*_, *f*_*cat*_, *V*, and *V*_*gap*_ (Table 1) to simultaneously estimate *f*_*res*_ and *r*. These results are summarized in Table 1. Our inferred value of autocatalytic nucleation *r* = 2.1min^−1^ is comparable to previous estimates: 1.5 min^−1^ (Clausen and Ribbeck, 2007) and 1 min^−1^ (Petry et al., 2013) in meiotic egg extract supplemented with RanGTP. In the alternative scenarios, where MCAK-Q710 decreases the catastrophe rate or increases the depolymerization rate, our estimates of *r* and *f*_*res*_ are essentially the same (Table S2). Thus, our model recapitulates aster growth with reasonable parameter values and offers a new understanding for how asters grow to span large cytoplasms even when individual microtubules are unstable.

To date, few studies have rigorously compared the mechanistic consequences of plus-end-stimulated vs. polymer-stimulated nucleation. Above, we presented the theoretical predictions for aster growth by plus-end stimulated nucleation. In the SI, we also provide the results for polymer-stimulated nucleation including the critical nucleation rate Eq. S59. Both models of nucleation have qualitatively similar behavior including the gap velocity and recapitulate experimental observations of asters growing as traveling waves. Thus, in our case, the qualitative conclusions do not depend on the precise molecular mechanism of autocatalytic nucleation. In particular, the explosive transition characterized by the gap velocity is a general prediction of modeling microtubules as self-amplifying elements whose lifetime depends on their length.

By carefully defining and quantifying autocatalytic nucleation, future studies may be able to distinguish its precise mechanism. With plus-end-stimulated nucleation, the nucleation rate *r* has units of min^−1^ and describes the number of new microtubules generated per existing plus end per minute. With polymer-stimulated nucleation, the nucleation rate has units of *μ*m^−1^ min^−1^, and describes the number of new microtubules generated per micron of existing microtubule per minute. This difference has important implications for the structural mechanism of microtubule nucleation and for the prediction of cell-scale phenomena. For the issue of large aster growth, we propose specific experiments that might be able distinguish these scenarios (SI).

### Phase Diagram for Aster Growth

How do large cells control aster size during rapid divisions? We summarize our theoretical findings with a phase diagram for aster growth in Fig. 5. Small mitotic asters are represented by stationary asters found in the regime of bounded polymerization dynamics *J* < 0 and low nucleation rates. These model parameters must change as cells transition from mitosis to interphase to produce large growing asters. Polymerization dynamics becomes more favorable to elongation during interphase (Belmont et al., 1990; Verde et al., 1992). This may be accompanied by an increased autocatalytic nucleation of microtubules.

**Figure 5.**
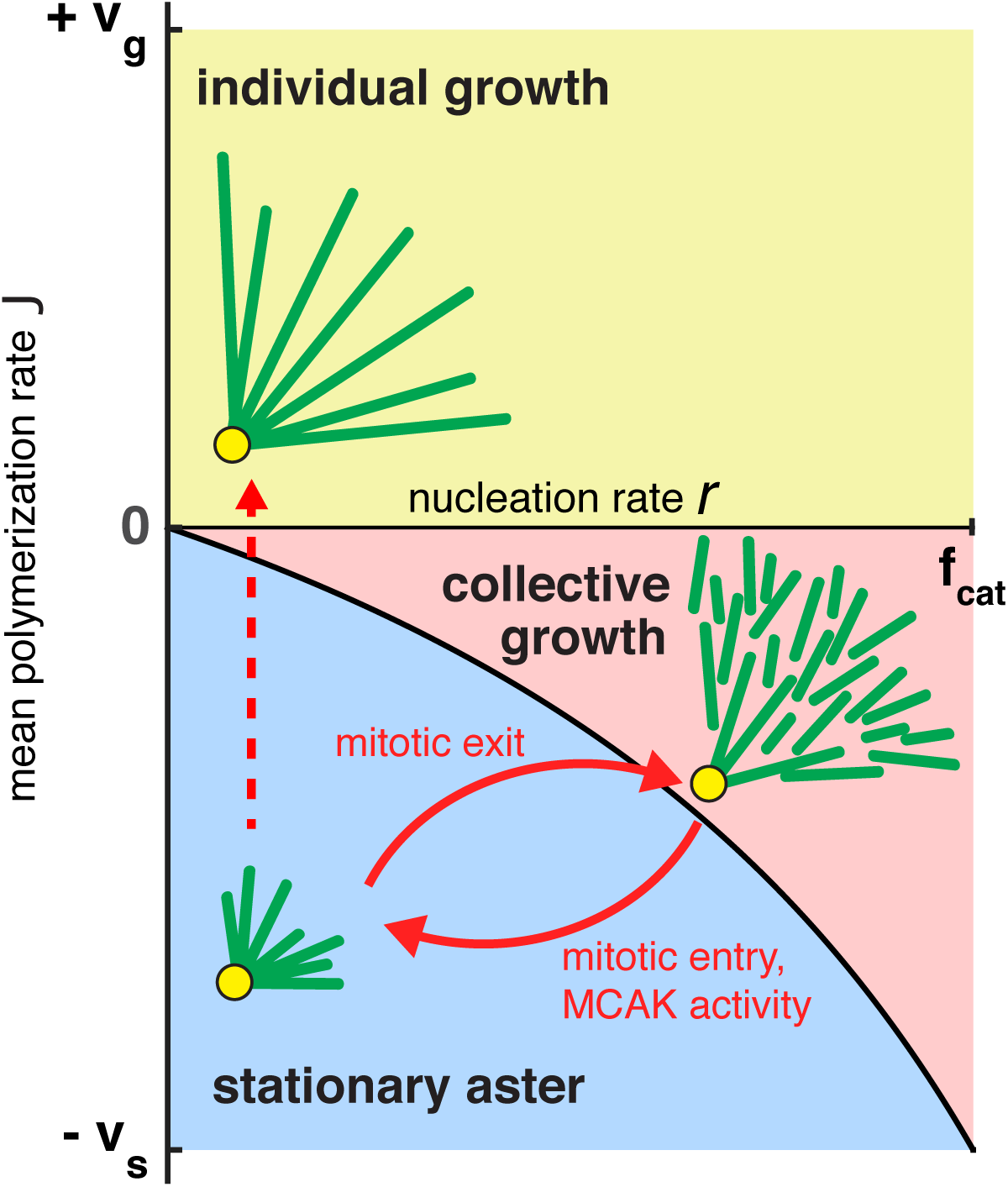
Phase diagram for aster growth. Aster morphology is determined by the balance of polymerization dynamics and autocatalytic nucleation. Small, stationary asters (*V* = 0), as observed during mitosis, occur at low nucleation *r* and net depolymerization of individual microtubules (*J* < 0). Net polymerization (*J* > 0) without nucleation (*r* = 0) produces asters that expand at rate *V* = *J* with dilution of microtubule density at the periphery and are thus inconsistent with experimental observations. The addition of nucleation to the individual growth regime changes these dynamics only marginally (yellow region); see SI. Alternatively, the transition from stationary to growing asters can be achieved by increasing the nucleation rate, *r*, while keeping *J* negative. Above the critical nucleation rate *r*_*c*_ starts the regime of collective growth (*V* as in Eq. (4), which is valid for *r*_*c*_ < *r* < *f*_*cat*_) that produces asters composed of relatively short microtubules (red region). The transition from stationary aster to collective growth may be achieved by crossing the curve at any location, but always involves an explosive jump in aster growth velocity, *V*_*gap*_. Reverse transition recapitulates the results of our experimental perturbation of MCAK activity (Fig. 4) and mitotic entry (solid arrows). We propose this unified biophysical picture as an explanation for the cell cycle dependent changes of aster morphology *in vivo*.

According to the standard model, increasing *J* to a positive value with no nucleation leads to asters in the “individual growth” regime. A previous study suggested the interphase cytoplasm is in the unbounded polymerization dynamics *J* > 0 (Verde et al., 1992), but our measurements of parameters used to calculate *J* differ greatly (Table 1). Individual growth regime is also inconsistent with the steady-state density of microtubules at the periphery of large asters in both fish and frog embryos (Ishihara et al., 2014a; Wühr et al., 2008, 2010). Experiments in egg extracts further confirm the addition of new microtubules during aster growth (Ishihara et al., 2014a) contrary to the predictions of the standard model. Furthermore, the presence of a high density of growing plus ends in the interior of growing asters in egg extract suggests that microtubules must be short compared to aster radius, and mean growth velocity must be negative, at least in the aster interior (Ishihara et al., 2014a).

By constructing a model that incorporates autocatalytic nucleation *r* > 0, we discovered a new regime, in which continuous aster growth is supported even when microtubules are unstable (*J* < 0). We call this the “collective growth” regime because individual microtubules are much shorter (estimated mean length of 20 *μ*m, Table 1) than the aster radius (hundreds of microns). Predictions of this model are fully confirmed by the biochemical perturbation via MCAK-Q710. The finite jump in the aster growth velocity (Fig. 4) is in sharp contrast to the prediction of the standard models of spatial growth (Fisher, 1937; Kolmogorov and Petrovskii, 1937; Skellam, 1951; van Saarloos, 2003). Spatial growth is typically modeled by reaction-diffusion processes that account for birth events and random motion, which, in the context of microtubules, correspond to the nucleation and dynamic instability of plus ends. Reaction-diffusion models, however, neglect internal dynamics of the agents such as the length of a microtubule. As a result, such models inevitably predict a continuous, gradual increase in the growth velocity as the model parameters are varied (Chang and Ferrell, 2013; Hallatschek and Korolev, 2009; Méndez et al., 2007; van Saarloos, 2003). The observation of finite velocity jump provides a strong support for our model and rules out a very wide class of models that reproduce the overall phenomenology of aster growth including the constant velocity and profile shape (Fig. 2). In particular, the observation of *V*_*gap*_ excludes the model that we previously proposed based on the analogy of aster growth and the Fisher-Kolmogorov equation (Ishihara et al., 2014b). The implications of *V*_*gap*_ for model selection are further discussed in SI.

### Collective Growth of Cytoskeletal Structures

Our theory allows for independent regulation of aster growth rate and microtubule density through the control of the nucleation rate and microtubule polymerization. Thus, cells have a lot of flexibility in optimizing aster properties and behavior. The existence of a gap velocity results in switch-like transition from quiescence to rapid growth and allows cells to drastically alter aster morphology with a small change of parameters. Importantly, the rapid growth does not require high microtubule density inside asters, which can be tuned independently.

Collective growth produces a meshwork of short microtubules with potentially desirable properties. First, the network is robust to microtubule severing or the spontaneous detachment from the centrosome. Second, the network can span arbitrarily large distances yet disassemble rapidly upon mitotic entry. Third, the structure, and therefore the mechanical properties, of the network do not depend on the distance from the centrosome. As a speculation, the physical interconnection of the microtubules may facilitate the transduction of mechanical forces across the cell in a way unattainable in the radial array predicted by the standard model (Tanimoto et al., 2016; Wühr et al., 2010).

The regime of collective growth parallels the assembly of other large cellular structures from short, interacting filaments (Pollard and Borisy, 2003) and is particularly reminiscent of how meiosis-II spindles self-assemble (Brugues and Needleman, 2014; Brugues et al., 2012; Burbank et al., 2007). Due to such dynamic architecture, spindles are known to have unique physical properties such as self-repair, fusion (Gatlin et al., 2009) and scaling (Good et al., 2013; Hazel et al., 2013; Wühr et al., 2008), which could allow for greater robustness and evolvability (Kirschner and Gerhart, 1998). Perhaps, collective growth is one of the most reliable ways for a cell to assemble cytoskeletal structures that exceed the typical length scales of individual filaments.

## MATERIALS AND METHODS

### Numerical Simulations

We implemented a finite difference method with fixed time steps to numerically solve the continuum model (Eq. (3)). Forward Euler's discretization scheme was used except exact solutions of advection equations was used to account for the gradient terms. Specifically, the plus end positions were simply shifted by +*v*_*g*_*δt* for growing microtubules and by −*v*_*s*_*δt* for shrinking microtubules. Nucleation added new growing microtubules of zero length at a position-dependent rate given by *Q*(*x*). The algorithm was implemented using MATLAB (Mathworks).

### Analytical Solution

We linearized Eq. (3) for small *C*_*g*_ and solved it using Laplace transforms in both space and time. The inverse Laplace transform was evaluated using the saddle point method (Bender and Orszag, 1999). We found the aster growth velocity as in Eq. (4). The details of this calculation are summarized in the Supporting Text (SI).

### Aster Growth Velocity Measurements

Interphase microtubule asters were reconstituted in *Xenopus* egg extract as described previously with use of p150-CC1 to inhibit dynein mediated microtubule sliding (Field et al., 2014; Ishihara et al., 2014a). Fluorescence microscopy was performed on a Nikon 90i upright microscope equipped with a Prior Proscan II motorized stage. EB1-mApple was imaged every 2 min with a 10x Plan Apo 0.45 N.A. or a 20x Plan Apo 0.75 N.A. objective. For the analysis of the aster growth front, a linear region originating from the center of asters was chosen (Figure 4-figure supplement 1). A low pass filter was applied to the fluorescence intensity profile and the half-max position, corresponding to the aster edge, was determined manually. The analysis was assisted by scripts written in ImageJ and MATLAB (Mathworks). Univariate scatter plots were generated with a template from (Weissgerber et al., 2015). EB1-mApple were purified as in (Petry et al., 2011), used at a final concentration of 100 nM. In some experiments, MCAK or MCAK-Q710-GFP (Moore and Wordeman, 2004) proteins were added to the reactions. Protein A Dynabeads coated with anti-Aurora kinase A antibody (Tsai and Zheng, 2005) or *Tetrahymena* pellicles were used as microtubule nucleating sites.

### Catastrophe Rate Measurements

Interphase asters were assembled as described above. Catastrophe rates and plus end polymerization rates were estimated from time lapse images of EB1 comets that localize to growing plus ends (Matov et al., 2010). The distributions of EB1 track durations were fitted to an exponential function to estimate the catastrophe rate. Spinning disc confocal microscopy was performed on a Nikon Ti motorized inverted microscope equipped with Perfect Focus, a Prior Proscan II motorized stage, Yokagawa CSU-X1 spinning disk confocal with Spectral Applied Research Aurora Borealis modification, Spectral Applied Research LMM-5 laser merge module with AOTF controlled solid state lasers: 488nm (100mW), 561nm (100mW), and Hamamatsu ORCA-AG cooled CCD camera. EB1-mApple was imaged every 2 sec with a 60x Plan Apo 1.40 N.A. objective with 2x2 binning. EB1 tracks were analyzed with PlusTipTracker (Applegate et al., 2011).

## Author Contributions

KI and KK developed and analyzed the model. KI performed the experiments and analyzed the data. KI, KK, and TM designed the research and wrote the manuscript.

## Acknowledgments

We thank the members of Mitchison and Korolev groups for helpful discussion. This work was supported by NIH grant GM39565 and by MBL summer fellowships. The computations in this paper were run on the Odyssey cluster supported by the FAS Division of Science, Research Computing Group at Harvard University. We thank Nikon Imaging Center at Harvard Medical School, and Nikon Inc. at Marine Biological Laboratory for microscopy support. We thank Ryoma Ohi for providing MCAK and expert advice. We thank Linda Wordeman for providing MCAK-Q710-GFP and expert advice. KK was supported by a start up fund from Boston University and by a grant from the Simons Foundation #409704. KI was supported by the Honjo International Scholarship Foundation.

